# Transcriptomic response to pyrethroid treatment in closely related bed bug strains varying in resistance

**DOI:** 10.1101/2024.03.15.584589

**Authors:** Chloé Haberkorn, Zaïnab Belgaidi, Romain Lasseur, Fabrice Vavre, Julien Varaldi

## Abstract

The common bed bug, *Cimex lectularius*, is one of the main human parasites. The world-wide resurgence of this pest is mainly due to globalization, and the spread of insecticide resistance. A few studies have compared the transcriptomes of susceptible and resistant strains. However, these studies usually relied on strains originating fromdistant locations, possibly explaining their extended candidate gene lists. Here, we compared the transcriptomes of two strains originating from the same location and showing low overall genetic differentiation (*F_ST_* =0.018) but varying in their susceptibility to pyrethroids, before and after insecticide exposure. In sharp contrast with previous studies, only 24 genes showing constitutive differential expression between the strains were identified. Interestingly, most of the genes with increased expression in the resistant strain encoded cuticular proteins. However, those changes were not associated with significant difference in cuticular thickness, suggesting that they might be involved in qualitative changes in the cuticle. In contrast, insecticide exposure induced the expression of a multitude of genes, mostly involved in detoxification. Finally, our set of transcriptome candidate loci showed little overlap with a set of loci strongly genetically differentiated in a previous study using the same strains. Several hypothesis explaining this discrepancy are discussed.

## 1 INTRODUCTION

*Cimex lectularius*, also known as the common bed bug, is an obligate blood-feeding parasite that mostly feed on humans. Physical and psychological disorders induced in their human preys can range from allergic reactions (Alexander, 1984), to psychosis and paranoia (Goddard and Deshazo, 2009). The current resurgence of this species, starting late 90s (Potter, 2011), is likely to be due to globalization, together with a widespread second-hand market (Doggett et al., 2004) and growing insecticide resistance (Davies et al., 2012). Pyrethroids are the main insecticides used to fight this pest and are therefore suspected to be the main pressure selecting resistance (Romero et al., 2007).

Pyrethroid resistance in *Cimex lectularius* is thought to be due to several mechanisms, common in insects. Cuticular resistance can first impair insecticide penetration in the body. Studies based on RT-qPCR assays showed that cuticular protein genes, such as chitin synthase (CHS) or cuticle protein, had higher transcript levels in resistant bed bug strains (Mamidala et al., 2012; Koganemaru et al., 2013). Following penetration of the cuticular barrier, detoxification metabolism may be mobilized to degrade or excrete the insecticide. Indeed, an increased activity of several enzymes such as cytochromes P450s (Romero et al., 2009) and esterases (Lilly et al., 2016a) has been observed in pyrethroid resistant strains. In addition, administration of piperonyl butoxide (PBO), a primary inhibitor of some cytochrome P450 monooxygenases, was associated with a significant decrease in resistance, further suggesting their involvement in resistance (Romero et al., 2009). Increased expressions of glutathione-S-transferases (GSTs) were also detected in resistant juvenile bed bugs (Mamidala et al., 2011). Finally, mutations can affect genes involved in the nervous system functioning, and more specifically in the voltage-gated sodium channels (*VGSC*), targeted by pyrethroids (Dang et al., 2014, 2015; Akhoundi et al., 2015; Balvín and Booth, 2018). A single non-synonymous mutation can alter *VGSC* conformation and hinder pyrethroids binding, thus conferring knock-down resistance (*kdr* mutations). The *kdr* mutant L925I (leucine 925 to isoleucine) has been identified in most bed bug populations (88% of the 117 strains tested in USA in Zhu et al. 2010, 100% of individuals in France in Durand et al. 2012), although some bed bugs had an additional V419L mutation (valine 419 to leucine, 40.9% in Zhu et al. 2010) or, more rarely, V419L alone (2.7% in Zhu et al. 2010).

To provide insights into the genomic sequences underlying insecticide resistance, we recently contrasted allele frequencies between resistant and susceptible strains of *C. lectularius* through a DNA pool-sequencing approach (Haberkorn et al., 2023). A 6 Mb superlocus showing high genetic differentiation between resistant and susceptible strains was identified. This region was enriched for SNPs showing strong genetic differentiation between strains, and for the presence of structural variants. This genomic region also contains the major QTL involved in pyrethroid resistance, as identified by Fountain et al. (2016). Additionally, several much shorter peaks of genetic differentiation were identified throughout the genome. These data thus suggest that the 6 Mb superlocus and possibly other smaller regions are involved in insecticide resistance, although the mechanisms and exact loci responsible for resistance remain to be confirmed.

In this work, using the same strains as in our genomic study, we compared the transcriptome of susceptible and resistant bed bugs exposed or not to insecticides, in order to identify genes possibly involved in insecticide resistance, exploring both constitutive and plastic response. Transcriptomic approaches are particularly relevant for identifying genes involved in detoxification, since their efficiency is often related to their level of expression (Li et al., 2007). A few studies, all published before the first bed bug genome was released, have addressed this question (Adelman et al., 2011; Bai et al., 2011; Zhu et al., 2013). However, since they all used 454 technology, quantification of each transcript was not possible (because of the relatively low throughput of this technology). Instead, a small set of transcripts/genes was further investigated using qRT-PCR. Since then, to our knowledge, only one resistance study using RNA-seq was conducted, leading to the identification of a huge quantity of transcripts differentially expressed between resistant/susceptible strains (15540 out of 51492 expressed sequence tags (ESTs), Mamidala et al. 2012). Although these studies provided significant insights into mechanisms possibly involved in resistance in bed bugs, they all suffered from several caveats. First, all the comparative studies mentioned so far were conducted on strains having very different genetic backgrounds. Indeed, most of them used the Harlan strain (sampled in Fort Dix, NJ in 1973) as the susceptible reference strain, and compared it to strains sampled from distant locations (800km away for Columbus, OH in Mamidala et al. 2012 and Bai et al. 2011, and 440km for Richmond, VA in Adelman et al. 2011) which probably differ in many traits unrelated to resistance phenotype. Consequently, it is unclear whether the genes or transcripts identified in these studies are differentially expressed as a result of selection by insecticides or other selective factors, or as a result of non-adaptive genetic differentiation (due to genetic drift). This may explain why so many transcripts were detected as differentially expressed in Mamidala et al. (2012). Additionally, these studies focused on constitutive differences between strains, since no exposure to insecticide was performed. This may limit our power to detect important genes involved in resistance, since resistance genes are often induced upon insecticide exposure (Guedes et al., 2017; Poupardin et al., 2008).

In the present study, we analyzed the whole protein-coding transcriptome of two bed bugs strains differing in resistance to pyrethroids. The two strains, that were either exposed or not to pyrethroids, were genetically very similar, since the overall index of differentiation (*F_ST_*) was only 0.018 (Haberkorn et al., 2023). Expression levels were then compared in order to identify (i) genes showing constitutive differences between the two strains, (ii) genes whose expression is altered after pyrethroid exposure, and (iii) genes whose expression is differently altered after insecticide exposure depending on the strain (interaction term). Whereas category (iii) can highlight both constitutive difference between strains or plastic response, category (ii) shows a shared plastic response across strains. We then crossed these results with our previous whole genome analysis on the very same strains (Haberkorn et al., 2023), in order to test whether the set of candidates obtained through the genomic and transcriptomic datasets significantly overlapped. Since we enlightened several cuticular genes over-expressed in the resistant strain with numerous non-synonymous mutations in close proximity, we explored the possibility of a cuticular thickening in this strain, as it has been observed in other resistant strains (Lilly et al., 2016b).

## 2 MATERIALS AND METHODS

### 2.1 Insects

The two strains used in this study were provided by Cimex Store Ltd (Chepstow, United Kingdom). The susceptible strain, London Lab (collected in London, Great Britain), was collected before the massive use of insecticide and raised in laboratories for over than 40 years. On the contrary, London Field (collected in 2008 in London, in Great Britain) was moderately resistant to pyrethroids.

Insects were kept isolated before imaginal moult in 24-wells Petri dishes containing accordion-folded blotting papers, serving as harborage. Bed bugs were maintained at 25°C, 40% relative humidity (RH), and a photoperiod of 12:12h. Since males perform the so-called traumatic insemination to copulate, which can be costly for their partners (Stutt and Siva-Jothy, 2001), insecticide exposures were conducted only on 7-day-old unfed virgin adult females.

### 2.2 Topical Assay

Insects were treated with a discriminating dose of 1 ng/*µ*L of deltamethrin, as assumed to reflect the LD_50_ of Cimex-Store resistant strains (Haberkorn et al., 2023). Insecticidal assays were carried out with deltamethrin (98% purity, Cluzeau, Sainte-Foy-La-Grande, France), a pyrethroid, as it remains the most used insecticide family recently for bed bug control. Four replicates of 8 insects were performed both for control and 1 ng/*µ*L insecticide exposure. Bed bugs were previously immobilized by placing them in a Petri dish on ice for 5 minutes. Topical applications were then made onto the ventral surface of the thorax, between the coxae, with a 50-µL glass syringe attached to a repeating dispenser (Hamilton Co., Reno, NV). Treated insects were exposed to 1 µL of insecticide powder diluted in acetone, whereas control insects received 1 µL of acetone only.

Mortality was assessed after 24 h by flipping each insect on dorsal side with a featherweight forceps, to see if it was able to reverse on the ventral side (alive) or if its movements were not coordinated enough to do so (moribund, considered as dead). A fisher test was used to assess the significance of mortality difference between pairs of strains.

### 2.3 RNA extraction and sequencing

For each replicate, three individuals were randomly sampled out of the survivors for insecticide-exposed and control conditions, immediately flash-frozen in liquid nitrogen and stored at -80°C. Those individual extractions were realized for four replicates, up to a total of 24 bed bugs per strain. Bed bugs were extracted using the RNAeasy Mini Kit (Qiagen, Hilden, Germany) with Turbo DNAse treatment on column (Thermofisher, Waltham MASS, USA). RNA concentration was measured using Nanodrop, and Qubit with RNA HS Kit (Agilent, Santa Clara CA, USA).

Both reverse-transcription and sequencing were performed by Macrogen Europe (Amsterdam, Netherlands). Libraries were prepared using TruSeq stranded mRNA kit (Illumina, San Diego CA, USA), and sequencing was performed on an Illumina Novaseq6000 machine producing 40 million reads 2*100PE. The raw data have been submitted to the Sequence Read Archive (SRA) database of NCBI under BioProject PRJNA832557.

### 2.4 Transcripts quantification

Adapters have already been removed by Macrogen, and no trimming was performed considering the very high quality of the reads (Q30 >92%), assessed using Fastqc (Andrews, 2015). Reads were then mapped on the bed bug genome using the STAR v2.7.3a workflow (Dobin et al., 2013). We used the Clec_2.1 and associated GTF annotation file (GCF_000648675.2). Parameters used for the mapping were as follow: "sjdbOverhang" of 100 (read size - 1), a computed optimal "genomeSAindexNbases" of 13, "genomeChrBinNbits" of 18 and "quantMode" on "GeneCounts". This last option allowed STAR to count the number of reads per gene while mapping, using the GTF annotation provided above (file "ReadsPerGene.out.tab"). For all 48 samples sequences (24 samples per strain), the proportion of uniquely mapped reads was high (on average 84,09%, minimum 75,56%). Multiple mapping reads were discarded when over 10 occurrences (default parameter).

### 2.5 Mapping scaffolds on linkage groups

Fountain et al. (2016) constructed a genetic map for *C. lectularius*, using RAD-seq markers on an F2 recombined population. However, as the genome assembly of *C. lectularius* has since been updated (v1.0 to 2.1), we used Fountain’s RAD-seq data and R scripts to generate a genetic map based on the latest genome assembly, as described in Haberkorn et al. (2023). This allowed us to obtain 14 putative autosomes, called linkage groups (LG).

### 2.6 Differential Expression analysis on London strains

Reads counts by gene (except tRNA) were analyzed using the R package DESeq2 v1.34.0 (Love et al., 2014) with "lab" and "untreated" as the reference levels. A model with interaction between conditions treatment and strain was built. Only genes with at least 10 reads (summed over all samples) were kept, leading to a total of 12187 genes out of 13208 (total number of genes without tRNA).

Genes were considered as significantly differentially-expressed when the adjusted p-value was below 5% (*p_adj_* < 0.05) and when the fold change in expression was above 1.5, i.e Log_2_ fold change (LFC) > 0.58 (up) or < -0.58 (down), as in Nardini et al. (2012).

### 2.7 Functional analysis

In order to test whether some relevant biological functions were enriched in the set of candidate genes, we defined ten categories of genes potentially involved in resistance, similarly to Faucon et al. (2015). Six of them corresponded to metabolic resistance : "Binding/Sequestration" (6 genes), "GST" (14), "CCE" (39), "UDPGT" (39), "ABC transporter/MRP" (55), and "P450" (56). Other categories were separated as: "Other detox" (64), "Cuticle" (113), "Insecticide target and nervous system" (31), and "Redox homeostasis" (14 genes). Details about those 431 genes are provided in Haberkorn et al. 2023. Analysis of enrichment of candidate genes in each resistance category were performed by comparing their proportions in genomic regions of interest with whole genome, using R/stats default package (proportion test with FDR correction for multiple comparisons).

### 2.8 Associating differentially-expressed genes with outliers SNPs and structural variants

Outlier SNPs were obtained from our previous genomic analysis based on the intersection of three criterion : (i) the differentiation between the two strains originating from London should be high (top 5% *F_ST_* s), (ii) alleles should be in a derived state, if one considers that the alleles of the reference genome (the insecticide susceptible Harlan strain) are ancestral (at least for loci involved in insecticide resistance), (iii) (derived) alleles should be in higher frequency in London Field than in London Lab. With these criteria applied using only the two London strains, 100,941 outlier SNPs were detected (versus 576 in Haberkorn et al. (2023) that included another set of strains). Among these SNPs, 59,537 were distributed in 4,987 genes (out of the 12,187).

Using a combination of read-pair orientation, insert size information and read depth, the genomic analysis also revealed 650 inverted duplications, 509 tandem duplications and 1,338 inversions in London Field compared to the reference genome, with higher frequencies in London Field compared to London Lab. These putative structural variants were overlapping with 4,118 genes (Haberkorn et al., 2023).

Analyses of enrichment in differentially expressed (DE) genes across LG or scaffolds were performed using R/stats default package (binomial test with Bonferroni correction for multiple testing). We also checked whether DE genes had higher values of genetic differentiation, and the fold change value in constitutive DE genes whenever containing a structural variant or not, while performing t-tests with alternative "greater".

### 2.9 Preparation of cuticular samples for scanning electron microscope

Cuticular resistance, mentioned above, could be due to an increased thickness of the bed bug cuticle (Balabanidou et al., 2018). We measured this trait on ten control specimens (exposed with acetone only) of each strain picked up from the RNA-seq experiment. These samples were preserved dry at -80°C. The left middle leg of each specimen was separated from the body at the apical region of the femur in fresh PBS buffer. The next steps were performed by the "Centre Technologique des Microstructures" (CTMU) at University Lyon 1. Briefly, fixation was first carried out for 2 hours with 2% glutaraldehyde in 0.1M sodium cacodylate buffer. Samples were then rinsed three times for 15min in 0.2M sodium cacodylate buffer. Post-fixation was performed in 1% osmium tetroxide in 0.1M sodium cacodylate buffer. After a rapid rinsing with distilled water, gradual dehydration were made onto increasing baths from 30% to 100% alcohol (30, 50, 70, 80, 95, 100) for 10 min each, followed by two 15min baths in propylene oxide. Progressive impregnation with EPON resin were then performed, followed by polymerization at 56°C (i.e. resin hardening in an oven). Blocks were then surfaced on the middle part of each tibia at room temperature with a diamond knife, and copper-plated with a 10nm deposit.

### 2.10 Electron microscope imaging and image analysis

Observations were made by CTMU laboratory with a Zeiss Merlin VP Compact SEM at 3 kV. Each sample was individually observed and a picture was captured with a working distance of 5mm, scan speed of 9, aperture size of 60 µm, and magnification of 729X.

We then analyzed raw images using Fiji in ImageJ2 (version 2.9.0/1.53t) (Rueden et al., 2017). The scale given in microns on the raw image was converted into a number of pixels to measure the cuticle thickness. A circle pattern with 12 radii was affixed to each image, enabling measurements to be taken from the point where the circle intersects the outside of the cuticle to the inside of the cuticle perpendicularly, whilst avoiding obvious debris, damage, or setae as made by Lilly et al. (2016b). Hence, 12 measurements were made for each cuticular cross-section sample. The experimenter was blind to the resistance status of the sample to avoid any bias. Cuticle thickness was compared between London Lab and London Field strains in a repeated measures ANOVA framework.

## 3 RESULTS

### 3.1 Assessment of resistance or susceptible phenotype

London Lab and London Field strains were first exposed to deltamethrin (1 ng per insect), in order to assess their pyrethroid resistance status. This dose was chosen based on a previous analysis showing significant differences in mortality between the strains (see Fig. 1 in Haberkorn et al. (2023)). As expected, the mean mortality was significantly higher for London Lab compared to London Field (59.4% versus 31.3%, n=24 for each strain, fisher test with p-value=0.044), although the difference was expected to be greater.

**FIGURE 1.**
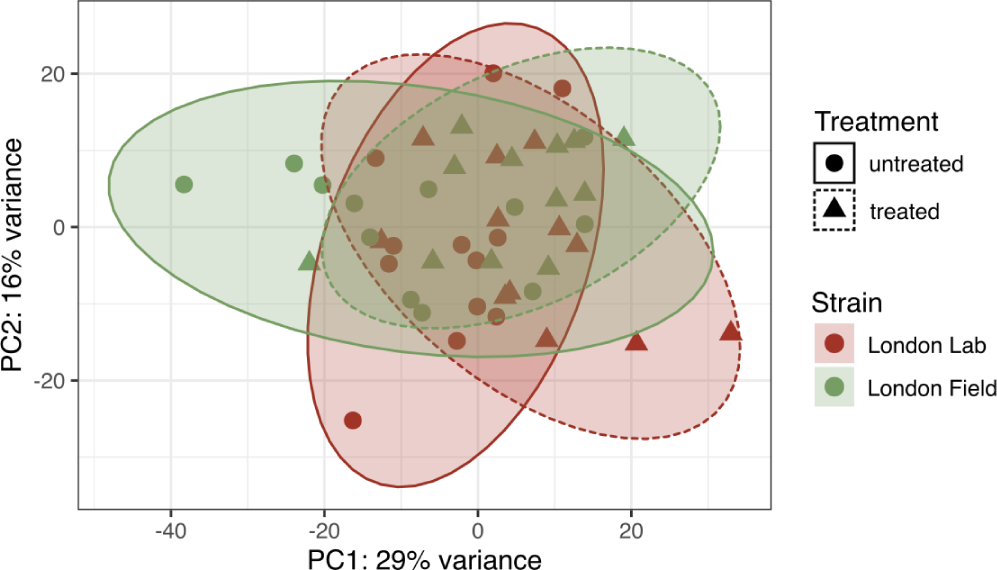
Projection of London Lab and London Field populations on the top two principal components using PCA. Read counts obtained from DESeq were processed with *vst*, which computes a variance stabilizing transformation. Individuals are separated within a population between survivors of insecticide treatment and untreated. Each point represents the top 500 most variable genes of a sample (default parameter for plotPCA function).

### 3.2 Principal component analysis does not reveal differences in expression profiles between resistant and susceptible strains

RNA-seq was performed in order to compare the transcriptome of the two strains both in the absence of insecticide (controls) and after insecticide exposure. The read counts table obtained after RNA-seq was first analyzed by Principal Component Analysis (PCA) (Figure 1). The sum of these two axes explained 45% of the variance (53% by adding PC3). No clear pattern was observed, neither between strains, nor between treated and untreated samples. This suggests that the transcriptomes were overall relatively similar between strains and between treated and untreated samples.

### 3.3 Analysis of differential gene expression

Variance in read counts was decomposed using a model including a strain effect (London Lab and London Field), a treatment effect (treated and untreated) and their interaction. Genes were considered as differentially expressed when the fold change exceeded 1.5 (in absolute value) and when *p_adj_* < 0.05.

#### 3.3.1 Constitutive differences between resistant and susceptible strains

The "strain" effect was first analyzed in order to identify genes showing constitutive differences in expression between the two strains. Out of 12187 genes, 20 genes were detected as up-regulated and 4 as down-regulated in London Field compared to London Lab (Figure 2).

**FIGURE 2.**
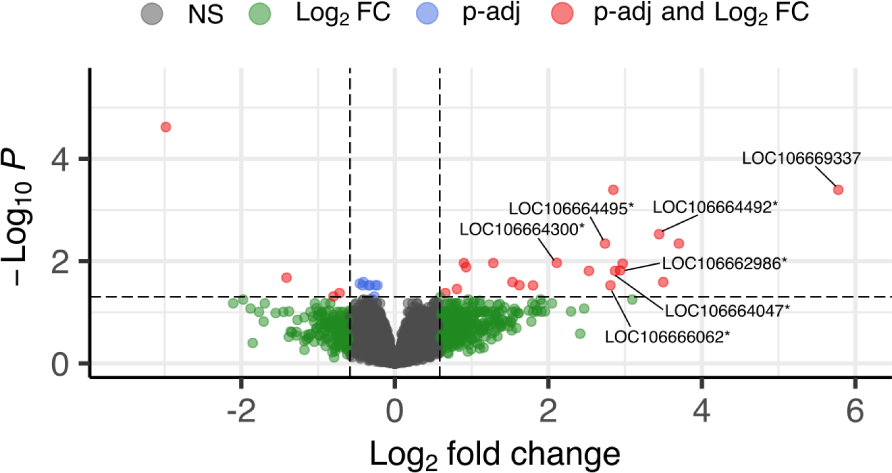
Volcano plot using enhanced volcano on strain effect (*n_genes_* = 12187, *p_adj_* < 0.05, LFC > or < |0.58|). Genes coding for cuticular proteins are labelled with a star.

Among the genes over-expressed in London Field, six were putatively involved in insecticide resistance which represents a highly significant over-representation for this category (proportion test, p-value = 5.13e-10). Strikingly, all six genes were coding for several cuticular proteins (Figure 3, Supplementary table 1), which also represents an over-representation of this specific category (p-value < 2.2e-16). Among over-expressed genes, the one showing simultaneously the highest LFC and lowest *p_adj_* encoded a putative leucine-rich repeat-containing protein DDB_G0290503 (106669337). To date, this gene is not known to be involved in insecticide resistance.

**FIGURE 3.**
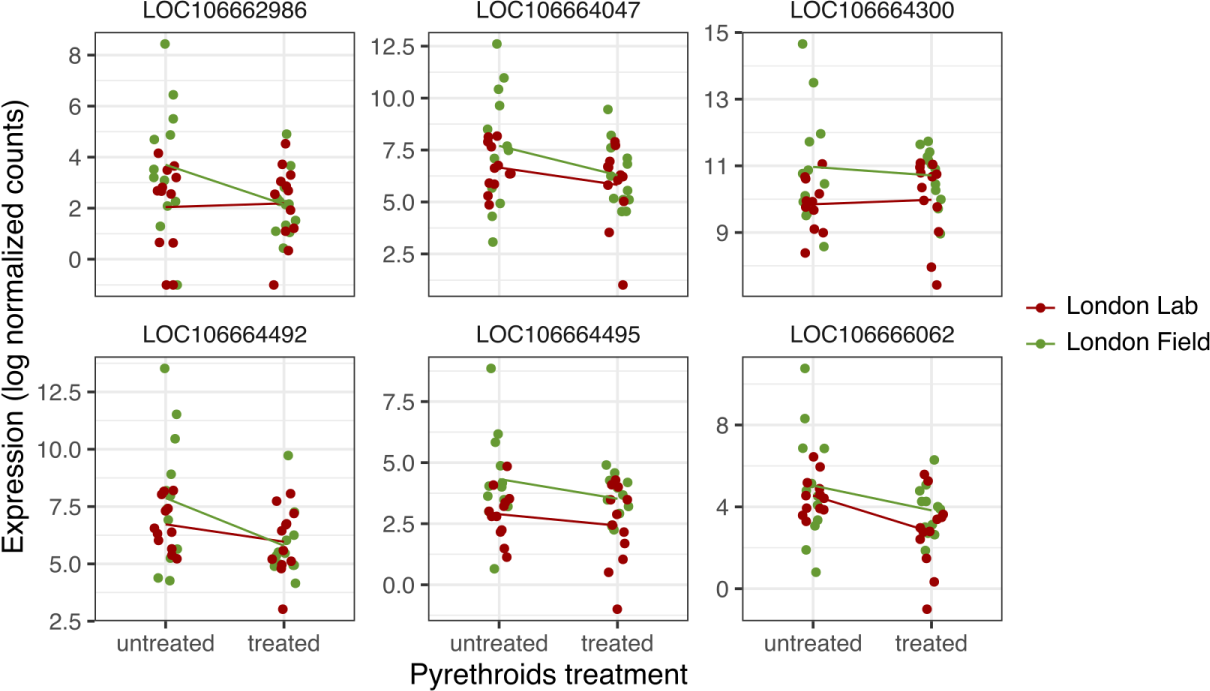
Cuticular proteins detected as significantly over-expressed in London Field. Expression is given in log2 of normalized count.

#### 3.3.2 Insecticide-altered genes

The "treatment" effect was then analyzed, i.e. the difference between untreated and treated survivors. Genes were similarly filtered on LFC and adjusted p-values, leading to respectively 375 genes significantly up-regulated and 388 down-regulated after insecticide exposure (Figure 4).

**FIGURE 4.**
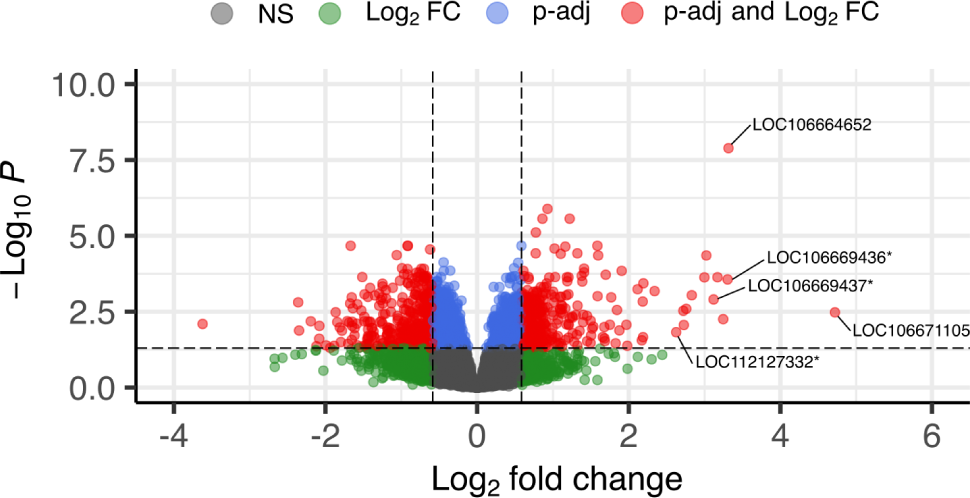
Volcano plot using enhanced volcano on treatment effect (*n_genes_* = 12187, *p_adj_* < 0.05, LFC > or < |0.58|). Genes coding for acetylcholinesterase-like proteins are labelled with a star.

Among the genes over-expressed in treated survivors, 20 were putatively involved in insecticide resistance (Figure 5, Supplementary table 2), which constitutes a significant enrichment (proportion test, p-value = 0.02). Two out of the six resistance categories among over-expressed genes were enriched, namely ABC transporters (*p_adj_* = 4.06E-03), and other detox (*p_adj_* = 8.19E-03).

**FIGURE 5.**
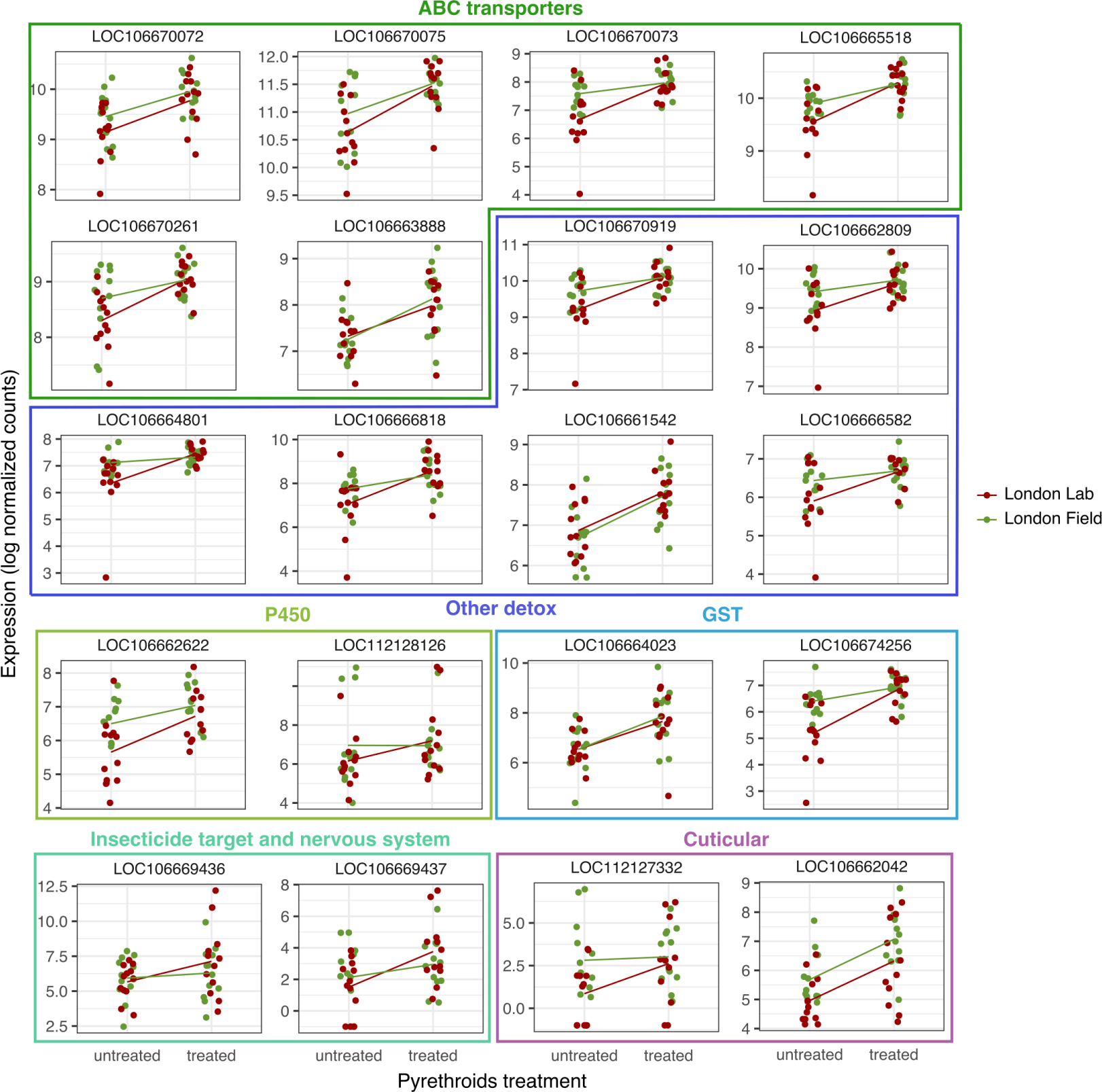
Resistance genes by category detected as significantly over-expressed in treated survivors. Expression is given in log2 of normalized count.

Most of the induced genes identified in a resistance category were involved in the detoxification metabolism: six ABC transporters, six various other detox (including transcription factor cap’n’collar and MAF, and three sulfotransferases), two GST and two P450. Three acetylcholinesterase-like and one cuticular protein were also detected. All acetylcholinesterase-like genes detected (labelled with a star on fig. 4) were in the top 15 of genes having the highest LFC. The up-regulated gene showing the highest LFC encoded a peroxidasin-like protein (106671105), whereas the up-regulated gene showing the lowest *p_adj_* encoded an elongation of very long chain fatty acids protein 7-like (106664652).

Of the down-regulated genes, 15 were detected in an insecticide resistance category (Supplementary table 3), which is not more than what is expected under the null hypothesis of no association between resistance and down-regulation (proportion test, p-value = 0.30). Similarly, no significant enrichment in any resistance gene category was detected (all *p_adj_* >0.05). Among the 15 genes identified, seven were coding for cuticular proteins, two for ABC transporters (both on LG 2), two for insecticide target and nervous system (one sodium channel protein Nach and one acetylcholinesterase-like), one from other detox category (sulfotransferase), two P450, and one Red/Ox gene (superoxide dismutase [Cu-Zn]-like). The down-regulated gene showing both the lowest LFC and lowest *p_adj_* was an uncharacterized protein (106673864).

#### 3.3.3 Interaction between treatment and strain

Only one gene showed a significant interaction between treatment and strain: a leucine-rich repeat protein soc-2: this gene had opposite expression patterns in the two strains upon insecticide exposure (106661213, Figure 6). Leucine rich repeat (LRR) motif are found in many immune receptors of animals, from *Homo* to *Drosophila* (Dushay and Eldon, 1998).

**FIGURE 6.**
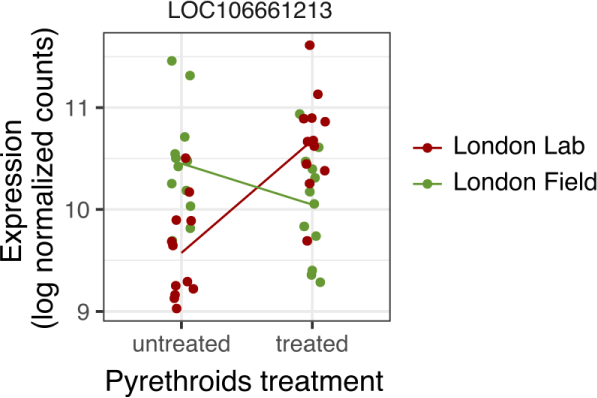
A gene putatively encoding a Leucine rich repeat (LRR) was associated with a significant interaction term between strain and treatment effects. Expression is given in log2 of normalized count.

### 3.4 Combining transcriptomic results with the genomic data

In a previous population genomic study conducted on the same two strains (Haberkorn et al., 2023), we identified a set of SNPs and structural variants (inversions, duplications) showing high between-strain genetic differentiation. These loci may therefore delineate genomic regions involved in the resistance phenotype. This genomic analysis revealed a clustering of candidate SNPs in some genomic regions, and particularly in 3 adjacent scaffolds constituting a 6 Mb region. This "superlocus" was furthermore significantly enriched in putative resistance genes, and in particular in *GSTs* and *P450s*, suggesting a clustering of those detoxification genes involved in insecticide resistance. One of the aims of the present study was to cross the population genomic analysis with the present transcriptomic analysis in order to identify some overlapping gene sets. Indeed, genes identified by both analyses are expected to be strong candidates for being involved in the phenotypic differences observed between the strains. In the following sections, we will therefore test whether the genes identified here are distributed evenly across the genome or whether they are clustered in some genomic regions, in particular in the highly differentiated genomic regions previously identified.

#### 3.4.1 Distribution of DE genes

We first tested whether the DE genes for treatment or strain effects were distributed evenly along the genome, at the linkage group (LG) or scaffold levels. Our hypothesis was that genes constitutively over-expressed in resistant individuals could be found in the superlocus previously identified. However, no pattern of enrichment was observed for differentially-expressed genes of both conditions, at any scale (binomial test with Bonferroni corrections, all p-values > 0.23). Hence, contrary to the SNPs that were clustered in some genomic locations, differentially-expressed genes for treatment or strain effects were scattered along the genome. Nonetheless, six significantly DE genes were identified in the 6 Mb superlocus, all of them being induced upon insecticide exposure. One of them was putatively involved in resistance, namely a GST on scaffold NW_019392763.1 (106664023, Supplementary table 2).

#### 3.4.2 Genetic differentiation of DE genes

*F_ST_* mean values were computed by gene, and compared between differentially-expressed genes and the non differentially-expressed pool. We hypothesized that *F_ST_* values should be higher for differentially-expressed genes, since both the genomic and the transcriptomic approaches are expected to be enriched for genes involved in insecticide resistance. Although no link was observed between *F_ST_* value and differential expression for the strain effect (t-test, p-value = 0.926), there was a marginally significant effect for the treatment effect (t-test, p-value = 0.059): *F_ST_* values were slightly higher for DE genes upon insecticide exposure compared to non-DE genes (*F_ST_* DE = 0.01356 > *F_ST_* non-DE = 0.00826). This suggests that DE genes upon insecticide exposure are located in genomic regions that are slightly more differentiated than the rest of the genome. However, the same test conducted within each individual LG did not reveal any pattern.

#### 3.4.3 Exploration of SNPs associated with transcriptomic differences between strains

Out of the 24 genes showing constitutive transcriptomic differences between LL and LF, twelve genes carried at least one outlier SNP, including four resistance genes - all being over-expressed cuticular genes (Table 1). Two of them carried outlier SNPs only in intergenic regions (106662986 and 106664047), whereas the others also had outlier SNPs in promoter (106664492) or intronic region (106666062). Finally, the only gene showing interaction between strain and treatment, i.e. the leucine-rich repeat protein soc-2 (106661213), was associated with five outlier SNPs (Table 2). Two of those SNPs were localized in intergenic regions, one in a promoter and two in introns.

**TABLE 1.** Outlier SNPs falling in resistance genes over-expressed in London Field when compared to London Lab strain. Outliers SNPs were determined as having their *F_ST_* value in top 5% quantile (i.e. empirical *pF_ST_* < 0.05) and the frequency of alternative allele (*F_al t_*) higher in London Field (LF) than London Lab (LL), according to our previous population genomic analysis Haberkorn et al. (2023). The position of each SNP is given, together with the distance to the nearest gene (Dist.) and its gene ID and annotation, reference to alternative nucleotide (R>A) and the location of the SNP (promoter=P, intergenic=Inter, intron=I). When several locations were available for the same single position, they were all reported and separated by a pipe.

| Scaffold | Position | Gene ID | Dist. | Annotation | Location | R>A | $F_{altLL}$ | $F_{altLF}$ | $F_{ST}$ | $pF_{ST}$ |
| --- | --- | --- | --- | --- | --- | --- | --- | --- | --- | --- |
| NW_019392649.1 | 1018810 | 106664492 | 17 | larval cut. A3A | P \| Inter | A>G | 0.22 | 0.64 | 0.25 | 0.044 |
|  | 1019770 |  | 977 |  | P \| Inter | C>T | 0.08 | 0.82 | 0.67 | 4.36E-04 |
| NW_019392685.1 | 759447 | 106662986 | 1832 | cuticle prot. 12.5 | Inter | T>C | 0.39 | 0.89 | 0.35 | 0.017 |
| NW_019392920.1 | 61585 | 106664047 | 15825 | larval cut. A1A | Inter | A>G | 0.45 | 0.88 | 0.29 | 0.032 |
|  | 62751 |  | 14659 |  | Inter | G>A | 0.73 | 1 | 0.31 | 0.027 |
|  | 67008 |  | 10402 |  | Inter | A>G | 0.64 | 1 | 0.27 | 0.037 |
|  | 68629 |  | 8781 |  | Inter | A>T | 0.46 | 0.87 | 0.27 | 0.037 |
| NW_019392993.1 | 2430659 | 106666062 | 7112 | cuticle protein | Inter | A>C | 0.10 | 0.69 | 0.46 | 5.91E-03 |
|  | 2430749 |  | 7022 |  | Inter | C>T | 0.15 | 0.63 | 0.36 | 0.017 |
|  | 2430766 |  | 7005 |  | Inter | T>G | 0.17 | 0.59 | 0.27 | 0.039 |
|  | 2431463 |  | 6308 |  | Inter | C>T | 0.24 | 0.70 | 0.29 | 0.032 |
|  | 2431466 |  | 6305 |  | Inter | C>T | 0.19 | 0.71 | 0.37 | 0.015 |
|  | 2431520 |  | 6251 |  | Inter | A>G | 0.24 | 0.75 | 0.36 | 0.016 |
|  | 2431528 |  | 6243 |  | Inter | G>A | 0.27 | 0.76 | 0.33 | 0.021 |
|  | 2434684 |  | 3087 |  | Inter | A>T | 0.18 | 0.67 | 0.30 | 0.028 |
|  | 2434689 |  | 3082 |  | Inter | T>C | 0.20 | 0.70 | 0.31 | 0.025 |
|  | 2434733 |  | 3038 |  | Inter | T>G | 0.27 | 0.80 | 0.35 | 0.017 |
|  | 2435135 |  | 2636 |  | Inter | T>C | 0.27 | 0.73 | 0.27 | 0.037 |
|  | 2438872 |  | - |  | I | T>A | 0.15 | 0.80 | 0.56 | 1.85E-03 |
|  | 2438882 |  | - |  | I | G>A | 0.13 | 0.70 | 0.45 | 6.55E-03 |
|  | 2445654 |  | - |  | I | G>T | 0.23 | 0.68 | 0.29 | 0.031 |
|  | 2452655 |  | - |  | I | T>C | 0.06 | 0.64 | 0.50 | 3.79E-03 |

**TABLE 2.** Outlier SNPs falling in gene over-expressed for interaction between strain and treatment. Outliers SNPs were determined as having their *F_ST_* value in top 5% quantile (i.e. empirical *pF_ST_* < 0.05) and the frequency of alternative allele (*F_al t_*) higher in London Field (LF) than London Lab (LL), according to our previous population genomic analysis Haberkorn et al. (2023). The position of each SNP is given, together with the distance to the nearest gene (Dist.) and its gene ID and annotation, reference to alternative nucleotide (R>A) and the location of the SNP (promoter=P, intergenic=Inter, intron=I). When several locations were available for the same single position, they were all reported and separated by a pipe.

| Scaffold | Position | Gene ID | Dist. | Annotation | Location | R>A | $F_{altLL}$ | $F_{altLF}$ | $F_{ST}$ | $pF_{ST}$ |
| --- | --- | --- | --- | --- | --- | --- | --- | --- | --- | --- |
| NW_019392653.1 | 1288539 | 106661213 | 1213 | LRR soc-2 | Inter | A>C | 0.29 | 0.89 | 0.50 | 3.92E-03 |
|  | 1288598 |  | 1154 |  | Inter | A>G | 0.38 | 0.81 | 0.26 | 0.042 |
|  | 1291912 |  | - |  | I \| P | C>A | 0.43 | 0.91 | 0.32 | 0.023 |
|  | 1299742 |  | - |  | I | G>C | 0.58 | 1 | 0.41 | 9.70E-03 |
|  | 1302774 |  | - |  | I | T>C | 0.39 | 0.83 | 0.30 | 0.030 |

#### 3.4.4 Exploration of structural variants associated with transcriptomic differences between strains

We first tested whether the genes identified in the present transcriptomic analysis showed a higher Log2 Fold Change value between strains when they contained a structural variant. The rationale was that structural variants may underlie transcriptomic changes, either through gene duplications or through rearrangements in promoter sequences.

Overall, this was not the case (t-test, p-value = 0.901). However, among the 20 over-expressed genes in London Field compared to London Lab, five were detected inside SVs (Supplementary table 4), including two cuticle protein A3A-like within a single putative inverted duplication located on LG5 (106664492 and 106664495). The three other genes were not part of any resistance category. Two genes out of the four under-expressed in LF compared to LL were part of SVs, but none of them had annotation suggesting their involvement in insecticide resistance. Finally, no structural variants were detected as overlapping with the only gene detected with an interaction between treatment and strain effects, namely LRR soc-2.

#### 3.4.5 Exploration of non-synonymous outlier SNPs associated with transcriptomic differences after insecticide exposure

Genes that are similarly induced upon insecticide treatment in both susceptible and resistant strains may still underlie variance in resistance if their coding sequences differ. We thus checked whether the DE genes for treatment (375 up-regulated, 388 down-regulated in LF) carried non-synonymous SNPs identified as outliers in our previous population genomic data. This analysis yielded 11 genes, including seven up-regulated and four down-regulated (Supplementary table 5). However, none of them belonged to a putative resistance category.

### 3.5 No significant difference in cuticular thickness between resistant and susceptible strains

As our transcriptomic analysis revealed a clear enrichment for genes encoding cuticle proteins among the constitutively differentially-expressed genes, we sought to compare cuticle thickness between the resistant and susceptible strains. Mean cuticular thickness were thus measured on bed bug legs extracted from both strains but no significant difference was observed (p=0.572, Figure 7).

**FIGURE 7.**
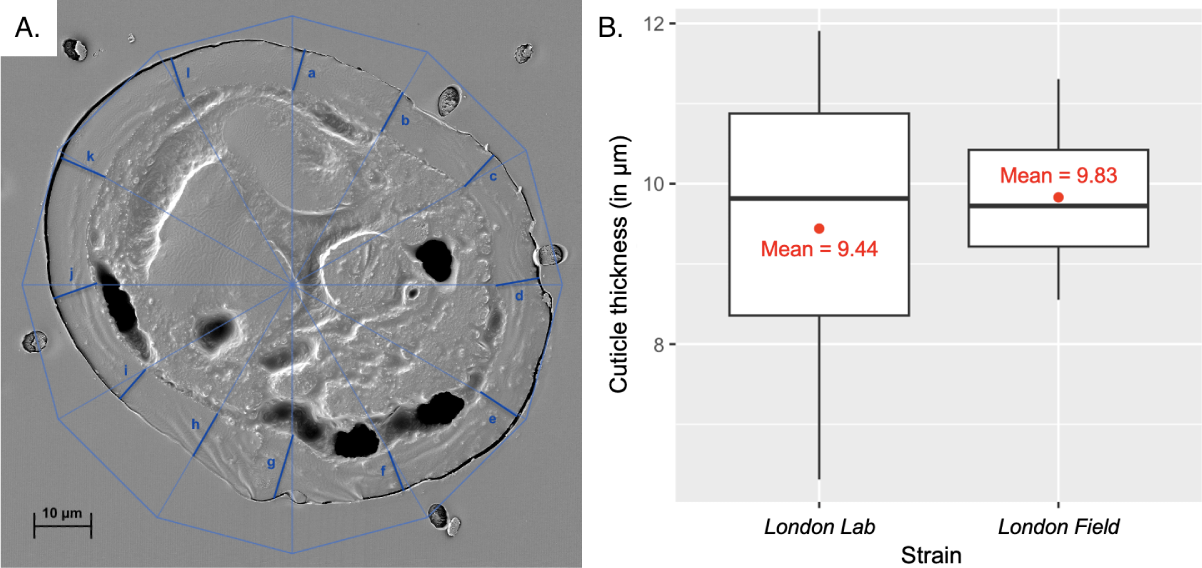
Analysis of *Cimex lectularius* cuticle. A. Example of a transverse section of a bed bug middle leg tibia in electron microscopy with the circle pattern (light blue) and subsequent 12 cuticle thickness measurements (blue, letters ’a’ to ’l’). B. Cuticle thickness (in µm) of London Lab (n=10) and London Field (n=10) strains. Mean values are represented with red dots.

## 4 DISCUSSION

The global resurgence of *C. lectularius* is likely caused by the evolution of insecticide resistance (Davies et al., 2012). A few transcriptomic analyses have been conducted on bed bugs, but only by comparing constitutive expression in resistant and susceptible strains with different genetic backgrounds (Zhu et al., 2012; Mamidala et al., 2012; Koganemaru et al., 2013). Consequently, it is unclear to what extent the genes or transcripts identified so far are truly relevant to insecticide resistance or other stresses, or whether they have any adaptive significance. Here, we provide a comprehensive transcriptomic analysis by comparing two bed bug strains with very similar genetic backgrounds (average genome-wide *F_ST_* = 0.018, Haberkorn et al. 2023) but differing in their resistance phenotypes. In addition to baseline constitutive gene expression, we analyzed gene expression before and after deltamethrin treatment in order to identify induced responses. We interpret our results in the light of a previous genome-wide analysis performed on the same strains, which identified genomic islands of exceptional differentiation (Haberkorn et al., 2023).

### 4.1 A small set of constitutively DE genes

We first compared the two strains in order to detect constitutive differences between strains that may underlie the observed difference in resistance. Only 24 genes (out of 12187) were detected as differentially-expressed, despite the insecticide assay confirmed the difference in resistance between the strains. However, the difference between LL and LF was lower than expected, which might be explained by individual variations in resistance level of the random insects selected.

Among the differentially-expressed genes, 20 genes were over-expressed in the resistant strain. This results is in sharp contrast with previous comparative whole transcriptome studies performed in bed bugs. Indeed, in Mamidala et al. (2012), several thousands of differentially-expressed transcripts were identified in the comparison between the two strains they used (Fort Dix, NJ and Columbus, OH). In Adelman et al. (2011) and Bai et al. (2011) which both relied on 454 sequencing followed by targeted RT-qPCR analysis, differences in transcripts abundance were detected for several categories of putative resistance genes: esterases, GSTs and several P450s. In contrast with these studies, a single category of putative resistance genes (cuticular genes) was identified in the small set of genes constitutively DE. Although part of the difference can be explained by the fact that they did not have a reference genome at the time (and thus had to rely on *de novo* assembly of the transcripts, which can inflate the number of inferred genes), another important explanation lies in the heterogeneity of the strains used. Indeed, in all three cases, the strains were collected in distant locations (>440km), meaning that they may differ in their allelic composition just because of other selective pressures experienced in the different environment they encountered (apart from insecticide), or because of genetic drift due to independent evolution. On the contrary, from our previous genetic analysis (Haberkorn et al., 2023), it appeared that London Lab and London Field strains were the most closely related of the four strains we analyzed with an overall low genetic differentiation (1.2% relative to the maximum possible value). This indicates that these strains are overall highly similar in their allelic composition. Genetic difference (that should underlie the constitutive expression differences observed) are thus more likely to be the consequence of selection for the major difference they have, i.e. the difference in pyrethroid resistance. This result thus challenges the view that transcriptomic changes involving various detoxification mechanisms (esterases, GSTs, P450) have evolved in pyrethroid-resistant bed bugs populations. However, it must be acknowledged that transcriptomic approaches such as the one carried out here as well as in the three previous studies mentioned above, are specifically designed to identify mechanisms that imply quantitative changes in effectors (i.e. typically in detoxification response). Consequently, these approaches do not allow the detection of other forms of resistance mechanisms involving qualitative rather than quantitative changes, such as target changes. It is thus still likely that the well characterized *kdr* mutation on *VGSC* plays a significant role in insecticide resistance in bed bug, as suggested by various studies (Zhu et al., 2010; Dang et al., 2015). Nevertheless, our result suggests that among the genes that experience quantitative transcriptomic changes, cuticular genes seem to play a predominant role. The cuticular barrier is the first encountered by a molecule in contact with a bed bug, and its construction is developed from the earliest larval stage, making any modification constitutive.

### 4.2 A key role for cuticular genes?

Six cuticular genes were detected as constitutively differentially-expressed, including three A3A larval cuticle proteins, one protein A1A and two uncharacterized cuticle proteins. Cuticular resistance has been observed in many pyrethroid-resistant insects, such as *Helicoverpa armigera* (Ahmad et al., 2006), *Anopheles gambiae* (Yahouédo et al., 2017) or *Culex pipiens* (Pan et al., 2009). In bed bugs also, several results also suggest that cuticle plays a central role in insecticide resistance (Balabanidou et al., 2018; Lilly et al., 2016b). Three transcripts encoding cuticular proteins (C2, C10 and C13) were detected as over-expressed in different resistant populations of bed-bugs (Zhu et al., 2013). Other studies showed that chitin synthase or larval/pupal cuticular protein had higher transcript level in pyrethroid resistant bed bug strains (Mamidala et al., 2012; Koganemaru et al., 2013). In particular, the larval cuticle protein A3A has also been detected among up-regulated transcripts in resistant insects (Gao et al., 2018; Zhang et al., 2022), including bed bugs (see Supplementary table 6 in Mamidala et al. 2012) . Interestingly, two A3A proteins detected as up-regulated in London Field strain were overlapping with a putative inverted duplication (Haberkorn et al., 2023). However, this inverted duplication was found both in London Lab and London Field strains, although in higher frequency in LF. Whether the small difference in the estimated frequencies for this inverted duplication (12.3% versus 5.7%) underlies the observed difference in A3A transcription is unclear but surely deserves additional investigations.

Cuticle-based resistance may arise thanks to an increase of its thickness, and or thanks to modifications of the composition of the cuticle Balabanidou et al. (2018). A previous morphological study showed a positive correlation between cuticle thickness and resistance status of bed bug strains (Lilly et al., 2016b). ’Resistant’ bed bug cuticles were significantly thicker than ’intolerant’ ones. In our present analysis, no significant difference was found when comparing leg cuticle thickness between resistant and susceptible strains (p = 0.572). Our data thus favours the scenario of a change in cuticle composition rather than an increased thickness. Future investigations should test more deeply this hypothesis.

### 4.3 Genetic variants associated with differences in expression between strains

Overall, the set of genes showing constitutive difference in expression was not particularly enriched for genetic variants identified by our previous population genomic analysis. However, a few outlier SNPs were detected, including within genomic regions annotated as promoters, that might be involved in the constitutive expression difference observed. These SNPs were part of the most differentiated loci detected in the whole genome analysis we conducted previously (Haberkorn et al., 2023). Four of these mutations were detected in introns of an over-expressed cuticular gene (106666062). This same gene was also associated with 11 outlier SNPs located in a small nearby flanking region (2636 bp to 7112 bp upstream). In one of the up-regulated larval cuticular A3A gene (106664492), two mutations were found in a flanking region annotated as promoter or intergenic sequence, depending on the transcript considered. Both flanking and intronic mutations may lead to transcriptomic changes, since they may serve as enhancers that ultimately recruit the RNA polymerase II on the promoter (Cannavò et al., 2016). For example, in *Nilaparvata lugens*, the up-regulation of the P450 genes CYP6AY1 and CYP6ER1 were associated with resistance to imidacloprid together with several mutations in their respective promoters. By site-directed mutagenesis, the authors were able to ascribe most of variance in up-regulation to a few set of single point mutations in the promotor sequences (Liang et al., 2018). The "resistant alleles" were associated with a higher binding activity for nuclear proteins, suggesting a higher recruitment of transcription factors (Liang et al., 2018). Thus, we may speculate that one or several of the mutations we highlight here are involved in the transcriptomic difference observed.

The unique gene showing an interaction between the strain and treatment effect encoded a leucine-rich repeat soc-2 protein. Although *soc-2* is not directly known to be involved in insecticide resistance, it seems to play a role in nicotinic acetylcholine receptor (nAChR) sensitivity, the target site of neonicotinoids (Gottschalk et al., 2005). High levels of neonicotinoid resistance have been observed in bed bug strains and attributed partially to enzymatic activities, without exploring the involvement of nAChR (Romero and Anderson, 2016). Although we did not test the resistance to neonicotinoids in the present study, it is reasonable to think that the recently collected LF strain (2008) has more chances of having been exposed to neonicotinoids in the past rather than the ancient LL strain (<1980) that was collected before the deployment of neonicotinoids (1990’s). Five SNPs showing high differentiation between LL and LF strains were detected in this gene, including one lying within a genomic region annotated as promoter. This polymorphism may thus be the genetic basis for the transcriptomic difference observed. More studies should be done to test the putative involvement of *soc-2* in insecticide resistance phenotype.

### 4.4 Genes induced upon insecticide exposure

Next, we aimed at identifying genes whose expression is modified upon insecticide exposure. In total, 375 genes were jointly induced in both strains. Among the 20 resistance genes up-regulated upon insecticide exposure, both ABC transporter and other detox categories of resistance genes were detected as significantly enriched (Supplementary table 2). Almost all of the resistance genes over-expressed upon insecticide exposure are detoxification enzymes, showing a strong generalist plastic response in both strains. Indeed, transcriptional plasticity of detoxification genes can allow to overcome insecticide application, even in non-resistant individuals (Mastrantonio et al., 2017). ABC transporters transcriptional plasticity in particular has previously been identified to enable adaptation of efflux capacity in *Tribolium castaneum* to get rid of insecticides (Rösner and Merzendorfer, 2020). This result also support the hypothesis that resistance genes have been acquired over millions of years of arms race to counter toxins produced by plants in a distant stinging-sucking past, still helping to excrete insecticides nowadays (Després et al., 2007).

The protein-coding genes which are similarly transcriptionally induced after insecticide exposure in both strains are not expected to explain a difference in resistance phenotype between the strains, unless they encode different proteins. Among the multitude of genes that are similarly induced in both resistant and susceptible strains, we identified a few showing this pattern (Supplementary table 5). Although none of them belonged to a resistance category, we considered that one deserved attention. A gene encoding a serpin B3 protein was associated with three nonsynonymous outlier SNPs, with the putative derived allele being fixed in the resistant strain. Additionally, this gene was associated in total with 28 non-synonymous mutations (including non-outlier SNPs) in our previous genomic analysis (Haberkorn et al., 2023). This gene was part of the top 1% of genes having the highest ratio between non-synonymous and synonymous mutations (*π_n_* /*π_s_* =4.67). Additionally, the gene was overall highly differentiated along its whole sequence, since the differentiation index computed at the scale of the whole gene was part of the top 5% (*F_ST_* =0.15). Overall, this strongly suggests that this gene has experienced positive selection in the recent past, leading to an increase in frequency of a putative derived allele. Although to our knowledge this gene has never been associated with pyrethroid resistance, the fact that it is also induced upon deltamethrin exposure renders its possible involvement in insecticide resistance plausible. The function of this particular gene is unclear, but serpin transcription has been shown to correlate with insecticide resistance in *Culex pipiens* mosquitoes (Li et al., 2016).

### 4.5 No evidence for differentially expressed genes to be clustered in a superlocus

Differentially-expressed genes, both between strains and between untreated and treated conditions, were dispersed across putative chromosomes (LG) and scaffolds of the bed bug assembly. More generally, there was no difference in average genetic differentiation between DE or non-DE genes. This is in sharp contrast with the high level of clustering of candidate SNPs observed in the bed bug genome, where a highly differentiated 6 Mb genomic region was identified (Haberkorn et al., 2023).

We consider three alternative explanations for this discrepancy: (i) most of the genomic regions identified previously are not associated with insecticide resistance, (ii) the resistance difference between the strains is mediated by quantitative or qualitative changes in messenger RNAs that were not measured in this study, or (iii) the resistance difference between the strains is not mediated by changes in mRNAs but rather by other mechanisms.

In support of hypothesis (i), although the two strains are highly comparable (*F_ST_* =0.018), we cannot exclude the possibility that part of the genetic differentiation observed is unrelated to insecticide resistance, i.e. related to other environmental factors, or even unrelated to adaptation, but rather due to genetic drift. Nonetheless, this does not exclude that some of the specific outlier SNPs we identified here (those located around or inside DE genes) are effectively underlying the transcriptomic changes detected as discussed above (Tables 1, 2, & Supp. table 5).

However, even if this is the case, we still expect a correlation between genetic differentiation and transcriptomic patterns, since even non-causal SNPs in linkage disequilibrium with causal SNPs should be statistically associated with the DE genes, leading to a signal of co-localization of constitutively-DE genes and regions of high genetic differentiation. This was not the case. A significant proportion of the candidate SNPs identified in our previous genomic analysis are therefore likely to be unrelated to insecticide resistance, i.e. related to other environmental factors or unrelated to adaptation, but but rather to drift.

In support of hypothesis (ii), we acknowledge that insecticide resistance may be triggered by between-strains differences in mRNA concentrations that are not quantified in this work. First, our transcriptomic analysis is focusing on adult insects, and any transcriptomic change relevant for insecticide resistance that would occur before adulthood would obviously not be detected. Indeed, previous studies have shown that the transcriptomic induction of some detoxification genes such as ABC transporter or P450 (Stevens et al., 2000), may confer insecticide resistance in the larval stages (Wang et al., 2023). Additionally, genes targeted by insecticide resistance such as *VGSC* may present non-synonymous changes underlying resistance, which are not accompanied by transcriptomic changes. They may be detected in the genomic differentiation analysis (this is the case for the *VGSC* locus in Haberkorn et al. (2023)) but surely not by the present transcriptomic analysis.

Finally, in support of (iii), a growing recent literature has brought to light examples of insecticide resistance mediated by non-coding RNAs that were not quantified in the present work. A recent study demonstrated that a long non-coding RNA (lncRNA) modulated the expression of a GST detoxification enzyme by competing with micro RNA (miRNA) binding, thus mediating cyflumetofen resistance (Feng et al., 2020). In *Aphis gossypii*, RT-qPCR and RNAi studies assessed the role of lncRNAs in acetyl-CoA carboxylase regulation, the target site of spirotetramat insecticide Peng et al. (2021). miRNAs are another type of ncRNAs which can affect insecticide resistance by themselves, as shown in *Spodoptera frugiperda* Yang et al. (2022). Indeed, injection of mi-RNAs-190-5p antagomir enhanced the expression of *CYP6K2*, improving insecticide tolerance. Hence, future studies should quantify the contribution of these mechanisms to the overall resistance pattern observed in *C. lectularius*.

## 5 CONCLUSION

In this study, most constitutive differences in expression between resistant and susceptible pyrethroid strains were due to cuticular genes. For four of those genes, we detected numerous outlier SNPs that may contribute to the differential of expression. After deltamethrin treatment, a majority of detoxification genes and especially ABC transporter G4 seemed to be at play for both strains, and without evident protein changes. These transcriptomic responses are thus unlikely to underlie the phenotypic difference observed between the strains, but rather of a common plastic response. Nevertheless, we identified a single gene (LRR soc-2) significantly associated with the interaction between our conditions, which is not yet known to be a resistance gene but whose role should be investigated. Little overlap was found between the genomic location of differentially expressed genes and the genomic regions showing high genetic differentiation. We considered three alternative hypotheses that may explain the pattern and should be disentangled in the future. To conclude, thanks to the combination of transcriptomics with previous genomics data, we identified a small set of genes (20 over-expressed in resistant strain, and one in interaction between resistance status and insecticide exposure) representing very strong candidates. Our rather stringent filters helped us to detect genes that, in our opinion, deserve to be functionally tested, to advance our understanding of previously unkown insecticide resistance mechanisms.

## acknowledgements

This work was performed using the computing facilities of the CC LBBE/PRABI. We thank the Symbiotron platform from the FR3728 BioEEnViS, especially Angelo Jacquet, and the Equipex+ InfectioTron (ANR-21-ESRE-0023) for facilities and equipment for rearing and experimentation on bed bugs. Electron microscopy studies have been done at the "Centre Technologique des Microstructures" - Claude Bernard University of Lyon, with special thanks to Lucie Geay. We also thank Dr Rike Stelkens (Stockholm University) for her gracious proofreading and valuable insights.

## data availability statement

The data that support the findings of this study are openly available on SRA under BioProject PRJNA832557.

