## Supplemental Tables 1-5 for "Transcriptomic response to pyrethroid treatment in closely related bed bug strains varying in resistance"

### 6 | SUPPLEMENTARY

| LG | Scaffold | Start | End | Gene ID | LFC | $p_{adj}$ | Annotation |
|---|---|---|---|---|---|---|---|
| 5 | NW_019392649.1 | 1017882 | 1018792 | 106664492 | 3.44 | 2.97E-03 | larval cuticle protein A3A |
| | | 1046176 | 1046953 | 106664495 | 2.74 | 4.53E-03 | larval cuticle protein A3A |
| 13 | NW_019392685.1 | 756768 | 757614 | 106662986 | 2.94 | 1.52E-02 | cuticle protein 12.5 |
| - | NW_019392920.1 | 77411 | 82689 | 106664047 | 2.87 | 1.55E-02 | larval cuticle protein A1A |
| - | NW_019392993.1 | 2437772 | 2454768 | 106666062 | 2.81 | 2.98E-02 | cuticle protein |
| - | NW_019393130.1 | 673262 | 674032 | 106664300 | 2.11 | 1.08E-02 | larval cuticle protein A3A |

**SUPPLEMENTARY TABLE 1** Resistance genes **over**-expressed in London Field when compared to London Lab strain for both untreated and treated survivors. LFC corresponds to the Log2 Fold Change between conditions (i.e. LFC of 3.44 means 10.85 fold change).

| LG | Scaffold | Start | End | Gene ID | LFC | $p_{adj}$ | Annotation |
|---|---|---|---|---|---|---|---|
| 4 | NW_019392661.1 | 361896 | 365391 | 106662622 | 0.90 | 2.60E-02 | cytochrome P450 6k1 |
| 5 | NW_019392644.1 | 647119 | 719373 | 106670072 | 0.62 | 1.03E-02 | ABC transporter G 4 |
| | | 749946 | 794199 | 106670075 | 0.78 | 1.88E-03 | ABC transporter G 4 |
| | | 1346254 | 1453427 | 106670073 | 0.99 | 3.20E-03 | ABC transporter G 4 |
| | NW_019392682.1 | 973424 | 1009568 | 106665518 | 0.66 | 6.15E-04 | ABC transporter G 4 |
| 9 | NW_019392638.1 | 815190 | 988569 | 106670919 | 0.79 | 1.54E-03 | segmentation protein cap'n'collar |
| | NW_019392675.1 | 595496 | 597255 | 106662042 | 1.77 | 1.07E-03 | cuticle protein 12.5 |
| 11 | NW_019392763.1 | 493269 | 495040 | 106664023 | 1.30 | 4.07E-03 | *GST* |
| - | NW_019392631.1 | 346739 | 535797 | 106662809 | 0.61 | 1.56E-02 | deacetylase/sulfotransferase |
| - | NW_019392668.1 | 644977 | 687162 | 106670261 | 0.69 | 3.65E-03 | ABC transporter G 1 |
| - | NW_019392709.1 | 60749 | 80839 | 106663888 | 0.67 | 2.70E-02 | ABC transporter G 23 |
| - | NW_019392742.1 | 1238897 | 1243972 | 106664801 | 0.88 | 4.89E-04 | protein-tyrosine sulfotransferase |
| | | 1771166 | 1773352 | 106666818 | 1.25 | 4.51E-03 | transcription factor MafB |
| - | NW_019392764.1 | 512150 | 523899 | 106661542 | 0.89 | 8.26E-03 | cytochrome b5-related protein |
| - | NW_019392803.1 | 7 | 886 | 112127332 | 2.63 | 1.50E-02 | *AChE* 1 |
| | | 7080 | 7993 | 106669437 | 3.12 | 1.25E-03 | *AChE* |
| - | NW_019392929.1 | 126420 | 138992 | 106674256 | 1.41 | 1.20E-04 | *GST* |
| - | NW_019393228.1 | 8 | 1146 | 112128126 | 1.88 | 1.31E-02 | probable cytochrome P450 9f2 |
| - | NW_019394179.1 | 823 | 3373 | 106669436 | 3.30 | 2.74E-04 | acetylcholinesterase |
| - | NW_019394202.1 | 1886643 | 1895153 | 106666582 | 0.60 | 2.86E-02 | carbohydrate sulfotransferase 11 |

**SUPPLEMENTARY TABLE 2** Resistance genes **over**-expressed in treated survivors when compared to untreated individuals for both London Lab and London Field strains. LFC corresponds to the Log2 Fold Change between conditions (i.e. LFC of 0.90 means 1.87 fold change).

| LG | Scaffold | Start | End | Gene ID | LFC | $p_{adj}$ | Annotation |
|---|---|---|---|---|---|---|---|
| 1 | NW_019392641.1 | 990484 | 1009113 | 106669934 | -1.25 | 7.83E-03 | SOD [Cu-Zn] |
| 2 | NW_019392653.1 | 1404658 | 1420128 | 106660996 | -0.81 | 1.09E-02 | ABC transporter G 23 |
| | | 1427987 | 1437993 | 106661276 | -0.80 | 4.20E-02 | ABC transporter G 23 |
| | NW_019392979.1 | 33885 | 34615 | 106665194 | -0.96 | 4.62E-02 | endocuticle structural glycoprot. |
| 4 | NW_019392730.1 | 1347727 | 1365982 | 106662446 | -1.41 | 5.20E-04 | sodium channel protein Nach |
| | NW_019392851.1 | 54557 | 58081 | 106670437 | -0.77 | 1.53E-02 | cuticle protein 18.7 |
| | | 220047 | 220929 | 106670525 | -1.24 | 3.87E-03 | pupal cuticle protein |
| | | 329822 | 331325 | 106670323 | -0.75 | 3.13E-02 | cuticle protein 18.7 |
| 6 | NW_019392719.1 | 1392866 | 1401379 | 106661029 | -0.96 | 2.62E-03 | pupal cuticle protein 20 |
| 13 | NW_019392637.1 | 3045820 | 3055263 | 106671932 | -0.80 | 3.18E-02 | *AChE* |
| | NW_019392781.1 | 529446 | 602890 | 106665955 | -0.74 | 2.84E-02 | sulfotransferase |
| - | NW_019392668.1 | 1276458 | 1279945 | 106670277 | -0.87 | 2.53E-02 | probable cytochrome P450 6a14 |
| - | NW_019392801.1 | 1599 | 16364 | 106669268 | -1.39 | 1.92E-02 | cytochrome P450 4C1 |
| - | NW_019392822.1 | 780550 | 784904 | 106665184 | -0.97 | 6.62E-03 | endocuticle structural glycoprot. |
| - | NW_019393130.1 | 625738 | 626515 | 106664854 | -0.85 | 2.03E-02 | larval cuticle protein A2B |

**SUPPLEMENTARY TABLE 3** Resistance genes **under**-expressed in treated survivors when compared to untreated individuals for both London Lab and London Field strains. LFC corresponds to the Log2 Fold Change between conditions (i.e. LFC of -1.25 means -2.37 fold change).

| LG | Scaffold | Gene ID | Annot. | TypeEvent | Size | $F$ LL | $F$ LF | $F_{ST}$ |
|---|---|---|---|---|---|---|---|---|
| 3 | NW_019392860.1 | LOC106674398 | uncharacterized | Inv | 15 kb | 0.10 | 0.21 | 0.022 |
| 5 | NW_019392649.1 | 106664492 | cuticle A3A | InvDup | 248 kb | 0.057 | 0.123 | 0.013 |
| | | 106664495 | cuticle A3A | | | 0.057 | 0.123 | 0.013 |
| 10 | NW_019392795.1 | 106667662 | GRP wall structural | InvDup | 1 M | 0.03 | 0.08 | 0.016 |
| | | | | InvDup | 241 kb | 0.16 | 0.17 | 1.18E-4 |
| | | | | Inv | 550 kb | 0.16 | 0.19 | 1.99E-03 |
| - | NW_019392939.1 | 112127884 | uncharacterized | Inv | 1 M | 0.07 | 0.09 | 1.02E-03 |

**SUPPLEMENTARY TABLE 4** Genes up-regulated in London Field when compared to London Lab individuals detected inside a structural variant (inversion "Inv", inverted duplication "InvDup" or tandem duplication "TanDup"). Size and frequencies of those variants in both strain were showed together with associated differentiation index ($F_{ST}$). Gene annotations have been shortened, but can be found using the gene ID on NCBI.

| DE | LG | Scaffold | Position | Gene ID | Annotation | Location | R>A | $F_{alt}LL$ | $F_{alt}LF$ | $F_{ST}$ | $pF_{ST}$ |
|---|---|---|---|---|---|---|---|---|---|---|---|
| up | 12 | NW_019392813.1 | 4400166 | 112127381 | uncharacterized | C \| I \| P | C>T | 0.20 | 0.68 | 0.32 | 0.023 |
| up | - | NW_019392656.1 | 2931030 | 106664583 | FAR wat | C \| I | T>G | 0.15 | 0.72 | 0.46 | 0.006 |
| | | | 2931057 | | | C \| I | C>A | 0.23 | 0.83 | 0.48 | 0.005 |
| | | | 2931088 | | | C \| I | G>T | 0.25 | 0.90 | 0.53 | 0.003 |
| | | | 2944507 | | | C | T>A | 0.10 | 0.52 | 0.27 | 0.036 |
| up | - | NW_019392664.1 | 1542428 | 106662188 | venom allergen | C \| I | T>G | 0.30 | 0.85 | 0.40 | 0.011 |
| | | | 1542429 | | | C \| I | T>G | 0.30 | 0.83 | 0.38 | 0.014 |
| up | - | NW_019392712.1 | 552763 | 106669984 | protein goliath | C | T>C | 0.06 | 0.50 | 0.30 | 0.030 |
| up | - | NW_019392775.1 | 745973 | 106663301 | AF4/FMR2 4 | C | A>G | 0.52 | 1 | 0.39 | 0.013 |
| up | - | NW_019392805.1 | 43356 | 106673166 | BCAT | C | G>C | 0.43 | 0.83 | 0.25 | 0.046 |
| up | - | NW_019392993.1 | 70657 | 106664949 | serpin B3 | C | A>T | 0.60 | 1 | 0.015 | |
| | | | 70664 | | | C | C>G | 0.55 | 1 | 0.37 | 0.015 |
| | | | 70665 | | | C | T>C | 0.55 | 1 | 0.015 | |
| do | 3 | NW_019392650.1 | 261464 | 106670335 | uncharacterized | C | A>T | 0.17 | 0.60 | 0.27 | 0.040 |
| do | 7 | NW_019392715.1 | 430046 | 106665593 | titin homolog | C | A>T | 0.48 | 0.93 | 0.30 | 0.029 |
| do | 8 | NW_019393011.1 | 184388 | 106665395 | solute carrier f26 m10 | C | C>A | 0.22 | 0.67 | 0.28 | 0.033 |
| do | - | NW_019392712.1 | 537273 | 106669983 | forkhead box P1 | C | T>C | 0.12 | 0.54 | 0.28 | 0.036 |

**SUPPLEMENTARY TABLE 5** Non-synonymous outlier SNPs in differentially expressed (DE) genes (up or down-regulated) for treatment effect. SNPs were selected with the frequency of alternative allele ($F_{alt}$) higher in London Field (LF) than London Lab (LL) and differentiation index $F_{ST}$ value in top 5% quantile (i.e. empirical $pF_{ST}$ < 0.05). The position of each SNP was gave, together with gene ID, gene annotation, reference to alternative nucleotide (R>A) and other putative location of the SNP whenever position was litigious due to the gap between transcripts annotation (coding=C, intron=I, promoter=P).
